# 64-Channel Carbon Fiber Electrode Arrays for Chronic Electrophysiology

**DOI:** 10.1101/697409

**Authors:** Grigori Guitchounts, David Cox

## Abstract

A chief goal in neuroscience is to understand how neuronal activity relates to behavior, perception, and cognition. However, monitoring neuronal activity over long periods of time is technically challenging, and limited, in part, by the invasive nature of recording tools. While electrodes allow for recording in freely-behaving animals, they tend to be bulky and stiff, causing damage to the tissue they are implanted in. One solution to this invasiveness problem may be probes that are small enough to fly under the immune system’s radar. Carbon fiber (CF) electrodes are thinner and more flexible than typical metal or silicon electrodes, but the arrays described in previous reports had low channel counts and required time-consuming manual assembly. Here we report the design of an expanded-channel-count carbon fiber electrode array (CFEA) as well as a method for fast preparation of the recording sites using acid etching and electroplating with PEDOT-TFB, and demonstrate the ability of the 64-channel CFEA to record from rat visual cortex. We include designs for interfacing the system with micro-drives or flex-PCB cables for recording from multiple brain regions, as well as a facilitated method for coating CFs with the insulator Parylene-C. High-channel-count CFEAs may thus be an alternative to traditional microwire-based electrodes and a practical tool for exploring the neural code.

## Introduction

In order to understand information processing in the brain, scientists must be able to take reliable measurements from the central nervous system (CNS). Ideal methods would allow us to record from all neurons in a brain (10^8^ in a rat or zebra finch^1^ at cellular spatial resolution and a temporal resolution on which neurons operate (kHz range) in behaving animals^2, 3^. While established methods for whole-brain monitoring (e.g. fMRI or PET) typically give poor spatial and temporal resolution, state of the art optical methods for imaging of genetically-encoded calcium indicators have yielded cellular spatial resolution while recording on the order of ∼10^5^ neurons simultaneously at low temporal resolution (∼0.5 Hz)^4, 5^.

In contrast to imaging methods, recording neuronal activity using electrodes provides spike-time temporal resolution but low cellular yield (state of the art simultaneously recorded neurons using Neuropixels probes is ∼3000 neurons using 8 probes, each of which contains 384 recording sites, yielding roughly one unit per recording site^6, 7^. A further limitation of electrode methods is their invasive nature: implanting large electrode arrays damages the tissue the electrodes are meant to record from^8, 9^, thus limiting the yield of recorded neurons, the longevity of the recording quality, and the ability to track the activity of individual neurons over long timescales. These are particularly pressing problems for investigations into how the neural code changes during learning, which unfolds over days, weeks or months^10–16^; or for the creation of practical brain-machine-interfaces (BMIs), which rely on the stability of recorded signals as well as the underlying neural code in order to decode a patient’s thoughts or intended actions^11, 17^.

Metal electrodes have been used for recording single neurons since the mid-20th century^18^. These are typically tungsten or PtIR wires, on the order of 12.5-50 µm in diameter^19, 20^. In rodent work, the typical approach involves arrays composed of individual metal wires spun into groups of four (tetrodes) and bundled into groups of up to 40 tetrodes, or 160 channels^21^. Monolithic silicon devices are a prominent alternative^6, 22, 23^ that allows for high-density recording sites and high channel counts at the expense of a large cross-section (e.g. 70 × 20*µm* in Neuropixels probes) and high stiffness, which make it a challenge to record on long-timescales (however, see Okun et. al^24^).

Carbon fiber electrode arrays (CFEAs) are an alternative to metal wires or silicon probes. Carbon fibers are thinner and more flexible than metal wires typically used, and produce a lower immune response after being implanted into the CNS^8, 25, 26^. We have previously demonstrated a method for producing multi-channel carbon fiber arrays^27^. However, the previous design was low-channel-count and required time-consuming individual preparation of the recording sites (each electrode’s tip had to be fire-sharpened to reduce impedance).

Here we demonstrate the design of 64-channel CFEAs and a method for bulk preparation of the recording sites. The tips were prepared by etching with sulfuric acid in order to increase surface area^28, 29^, followed by electroplating with PEDOT-TFB^30, 31^, resulting in a dramatically decreased impedance. Recordings in rat visual cortex demonstrate the feasibility of recording neural signals with this method. Designs for the 64-channel array are freely available online (https://github.com/guitchounts/electrodes), as are designs for flex-PCB-based 16-channel arrays, which allow for recording from multiple brain areas simultaneously. Finally, we share designs for laser-cut cartridges that facilitate preparation of CFs for insulation with Parylene-C.

## Results

### Assembly and tip preparation

In this report, we tested two types of CF arrays: a 16-channel version (assembly of which was reported in^27^, and a new, 64-channel version. For the latter, we designed a 3D-printed plastic block to hold the 64 CFs (Fig. 1a). As in the 16-channel version, carbon fibers were threaded through wells at the top of the block, and down through the exit holes on the bottom. The pictured design features four exit holes, arranged in a 2×2 grid, but this arrangement can be modified to an 8×2 grid of exit holes (design available on github.com) or a single exit hole to bundle the 64 CFs into one. After threading, fibers protrude from the top wells and from the bottom exit holes. To connectorize the array, the top-well-protruding wires are first de-insulated (by a flame passed swiftly over the wires, to burn off the Parylene-C), then pushed farther into the wells until they no longer protrude through the top. Then, a dab of silver paint is applied to the wells; once it spreads through the wells, excess paint is brushed off with a cotton-tipped applicator, and a connector is placed into the wells. In the 64-channel version, this is a 70-pin Hirose connector whose pins were folded down manually before insertion (Fig.fig:assemblya). The connector is then glued to the plastic block with super glue. The design is configurable to single-block 64-channel arrays (Fig. 1b) or four 16-channel arrays on flex-cable PCBs for multi-brain-region recordings (Fig.1c), and is compatible with micro-drive devices (Fig. 1d)^32^.

**Figure 1.**
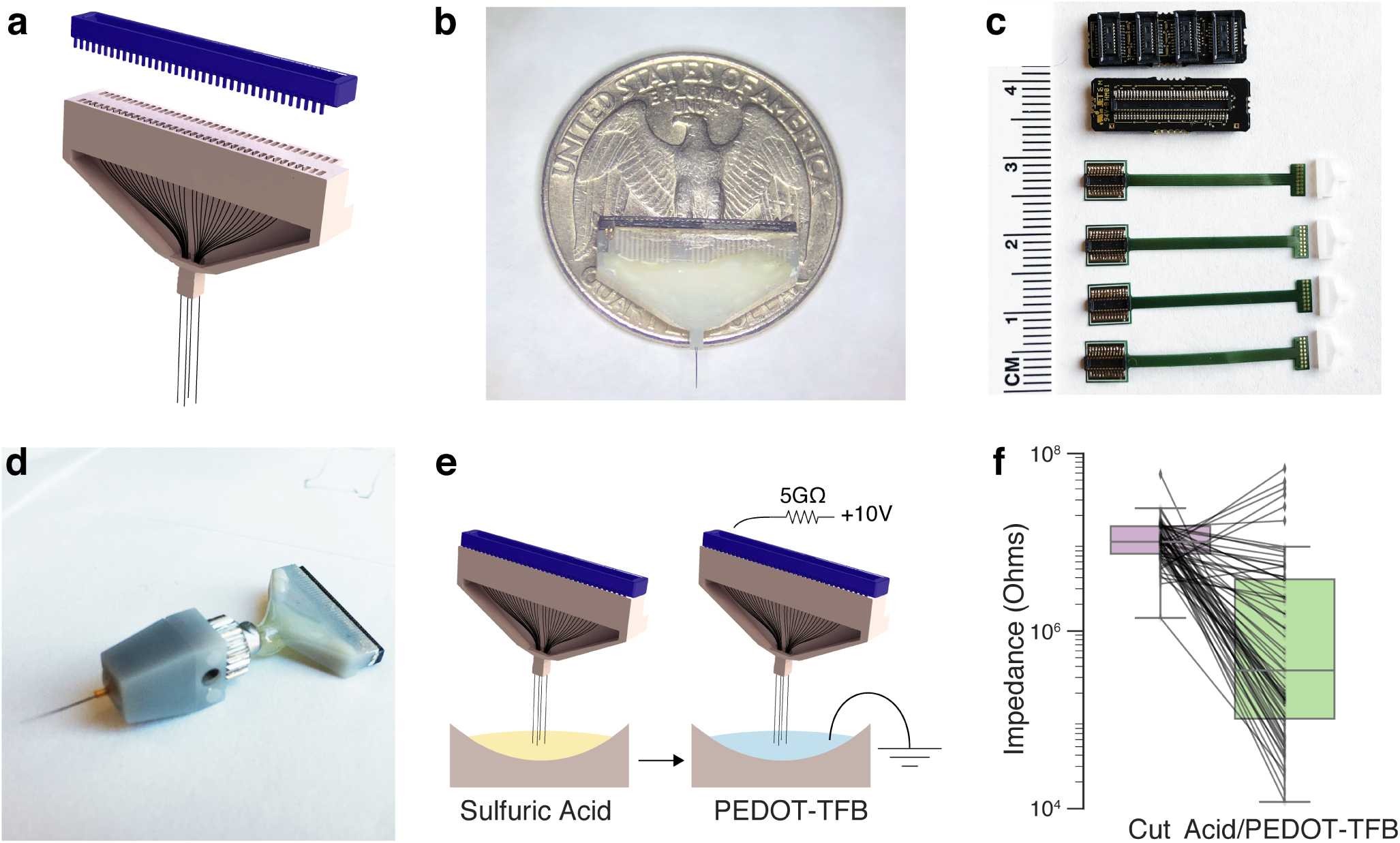
Assembly and Tip-Preparation. (**a**) Schematic of 64-channel 3D-printed block (grey) for holding carbon fibers (CFs). After threading the CFs through the block, the wells are filled with silver paint. A 70-pin Hirose connector (blue) mates with the wells. (**b**) Photo of assembled array with a US quarter. (**c**) System for multi-region recording using four 16-channel flex-PCB cables and 3D-printed blocks that mate with a PCB adapter for 64-channel recording (top). (**d**) A 64-channel CFEA mounted on a vented-screw-based micro-drive based on design in Anikeeva et. al^32^. (**e**) The recording sites are prepared by dipping the CF tips into 93% sulfuric acid for 5 minutes (left). After washing, the tips are electroplated with EDOT/TFB by passing a 2nA current for 1 minute per electrode. (**f**) Impedance of 64 wires on one array after cutting the tips with scissors (10.11 ± 0.94 MΩ, median ± s.e.m.) and after treatment with the acid and EDOT/TFB (0.36 ± 1.53 M Ω).

The impedance of untreated tips is too high for single-unit recordings in a fixed-implant preparation^33^, requiring treatment before recordings can be made. In a previous report, tips were prepared by individual fire-sharpening, which de-insulated ∼80 *µm* along the shaft of each wire, simultaneously sharpening it^27^. While successful, this technique is difficult to scale to high-channel-count arrays. To get around this problem, we turned to chemical treatment of the tips, first etching the tips using sulfuric acid in order to roughen the tip surface and increase the surface area^28, 29^, then electroplating the tips with PEDOT-TFB, which was previously shown to reduce impedance^30, 31^ (Fig. 1e). The resulting tips had significantly reduced impedances (before treatment: 10.11 ±0.94 MΩ median ±s.e.m. (*n* = 64 tips from one array), and 0.36 ±1.53 MΩ in the same tips after treatment with the acid and PEDOT-TFB (two-sided t-test on impedance before/after treatment of tips: *t* = 3.42, *p* = 0.001)) (Fig. 1f). We visualized the effect of this treatment on the tips using serial electron microscopy (Fig. 2).

**Figure 2.**
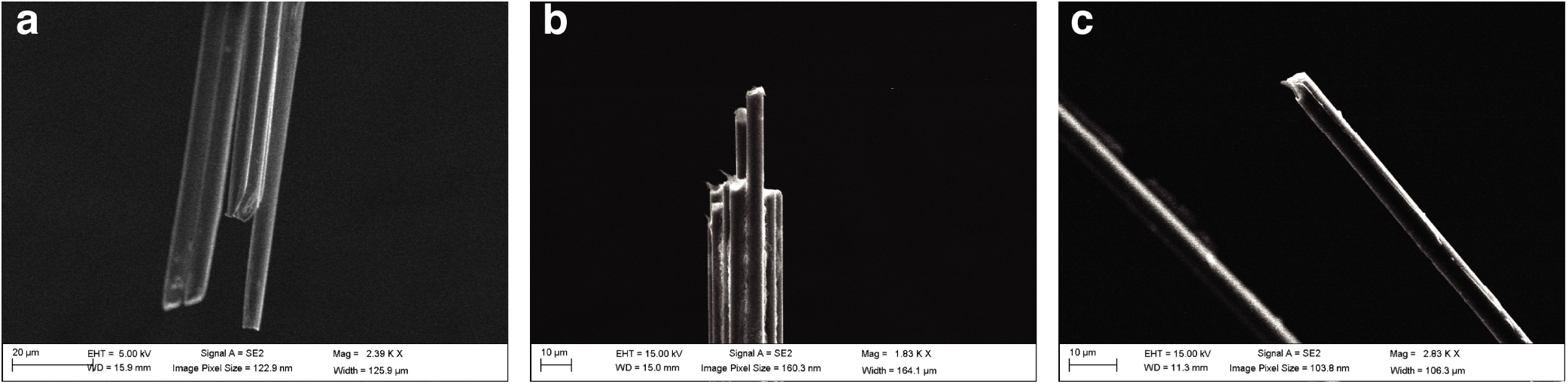
SEM Images of CF tips. (**a**) Bundle before treatment. (**b**) Bundle after acid treatment. (**c**) Close-up of a single fiber after electroplating with EDOT/TFB.

The CFEA require individual carbon fibers to be insulated with parylene-C before assembly, which in our previous design involved a painstaking manual procedure in which individual fibers were mounted on sticky notes and loaded into the parylene vapor deposition machine (PDS Labcoter 2010). To facilitate this process, we designed coating jigs (Fig. 3a-c) that hold multiple cartridges (Fig. 3d-e), each of which in turn hold several dozen carbon fibers splayed harp-like across two pieces of sticky notes. This design (Fig. 3f) can be laser-cut out of plastic and modified to various sizes of cartridges (to hold longer fibers, for example) or jigs (to hold more or fewer cartridges).

**Figure 3.**
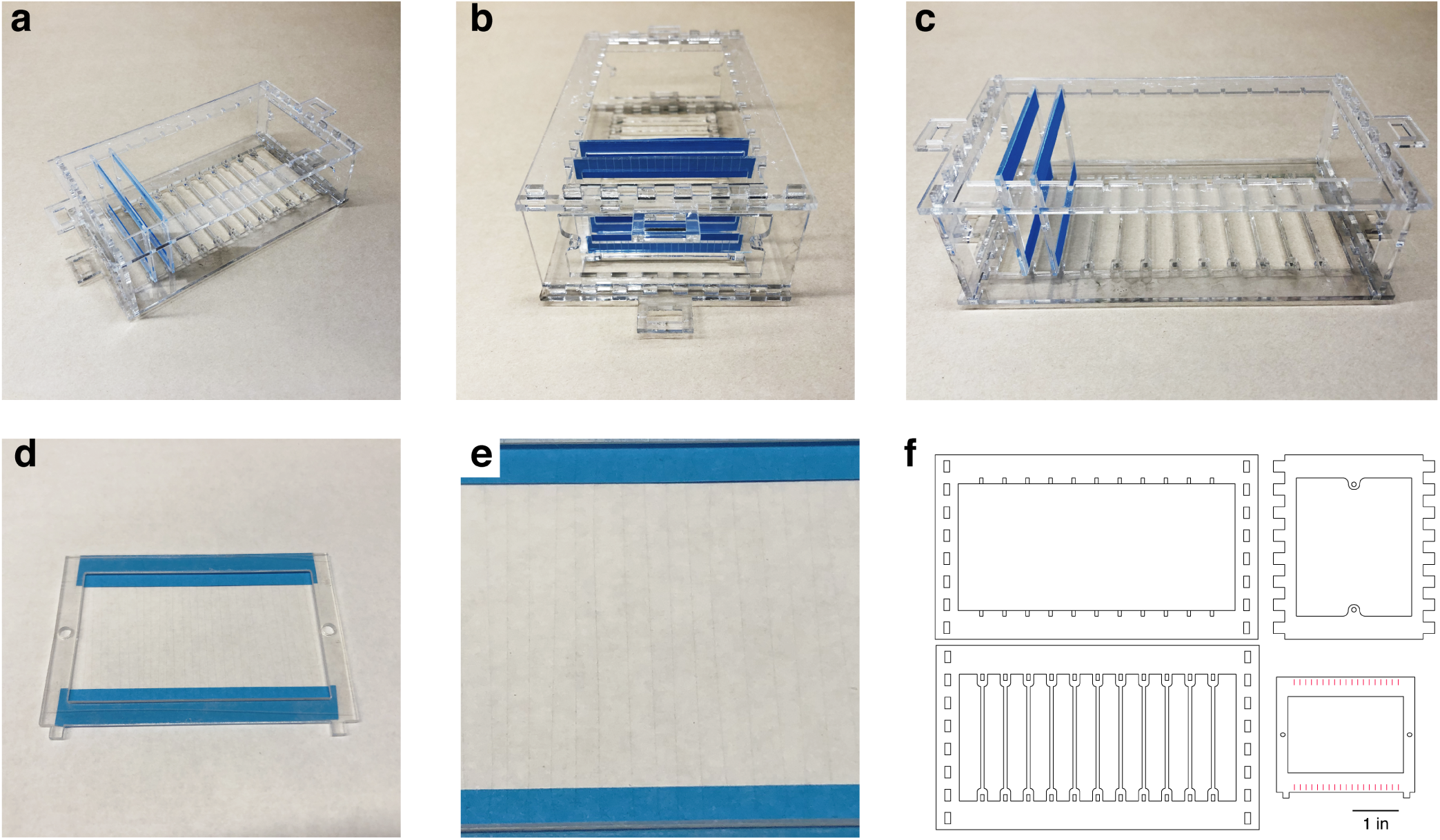
Parylene Coating Jig. (**a-c**) The assembled jig with two CF cartridges. (**d**) An example cartridge. Each cartridge holds 20-30 CFs, which are strung out across two sticky notes attached to the cartridge. (**e**) The fibers are placed across the cartridge like strings on a harp. (**f**) Jig and cartridge designs. Each jig holds 11 cartridges, but this design may be expanded as long as the jig fits into the Parylene deposition machine (PDS Labcoter 2010).

### Long-term recordings

We tested the longevity of acid/PEDOT-TFB-treated arrays by recording chronically from a 16-channel version of the array implanted in rat primary visual cortex (V1). Figure 4 shows examples from one such implanted rat. The left column shows two seconds of bandpass-filtered activity four days after the implant surgery (‘Day 4’), while the right column shows activity from the same array 55 days later. Spiking activity was prominent across most channels in both recordings, even though the noise increased over the two months.

**Figure 4.**
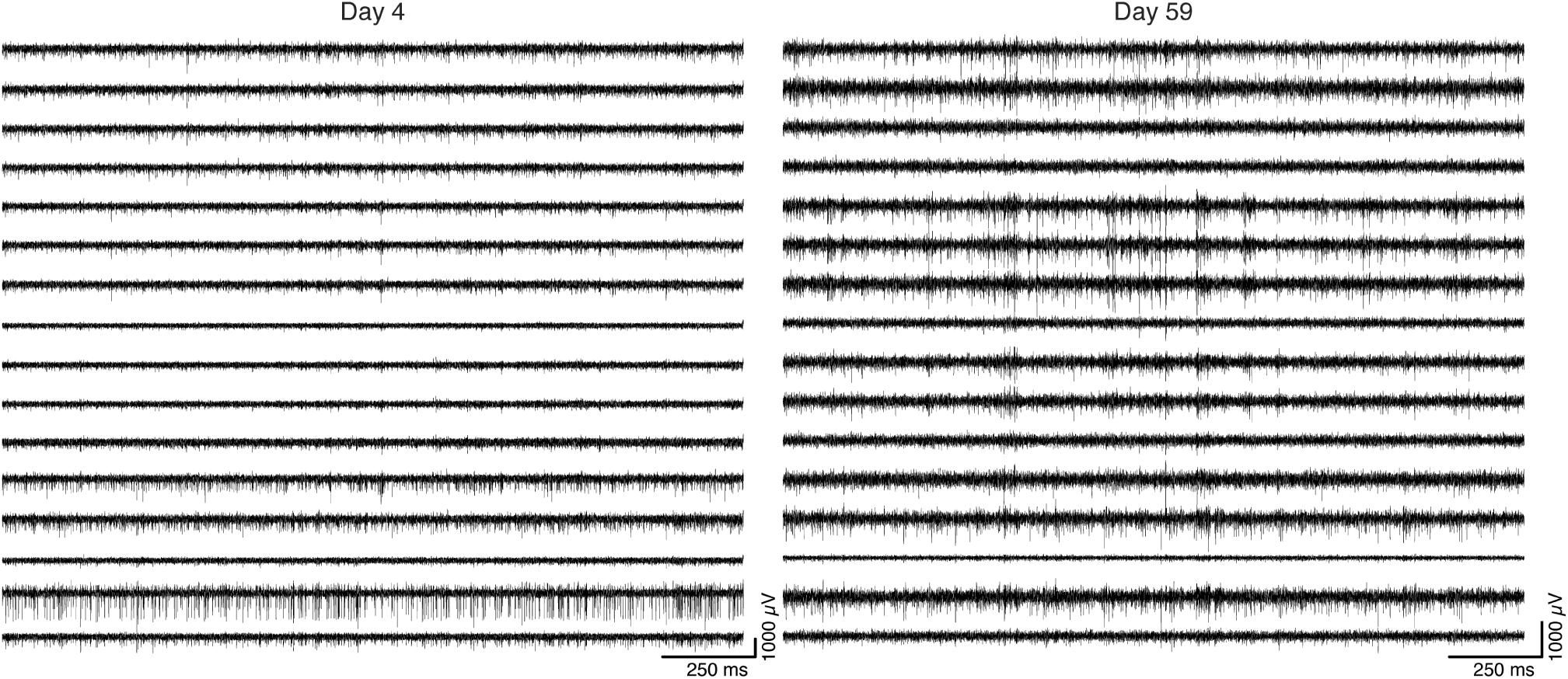
Example of 2 seconds of filtered activity traces from a 16-channel CF array chronically implanted in rat visual cortex. Left: activity sampled four days after implant surgery. Right: recording from the same animal’s array 55 days later.

### 64-Channel Recording and Responses to Visual Stimulation

We then proeeded to implant the 64-channel arrays into rat V1. Figure 5 shows example traces from one such recording while the animal was behaving freely in the recording chamber. Having determined that we could record spontaneous neuronal activity using chronically-implanted 64-channel CF arrays, we sought to determine whether these arrays could capture sensory evoked activity as well. For this, we presented a V1-implanted rat with a visual stimulus: flashed cage lights. The rat was free to behave in the recording chamber in the dark, while the cage lights were flashed (On for 500 milliseconds, Off for a uniformly random time between 400 and 600 milliseconds, for 225 trials.). This strong visual stimulus produced evoked multiunit (MU) responses (Fig. 6a,b) and LFPs (Fig. 6c). Figure 6a shows one trial (grey patch indicates time when lights were On); Figure 6b,c show trial-averaged MU and LFP responses, respectively.

**Figure 5.**
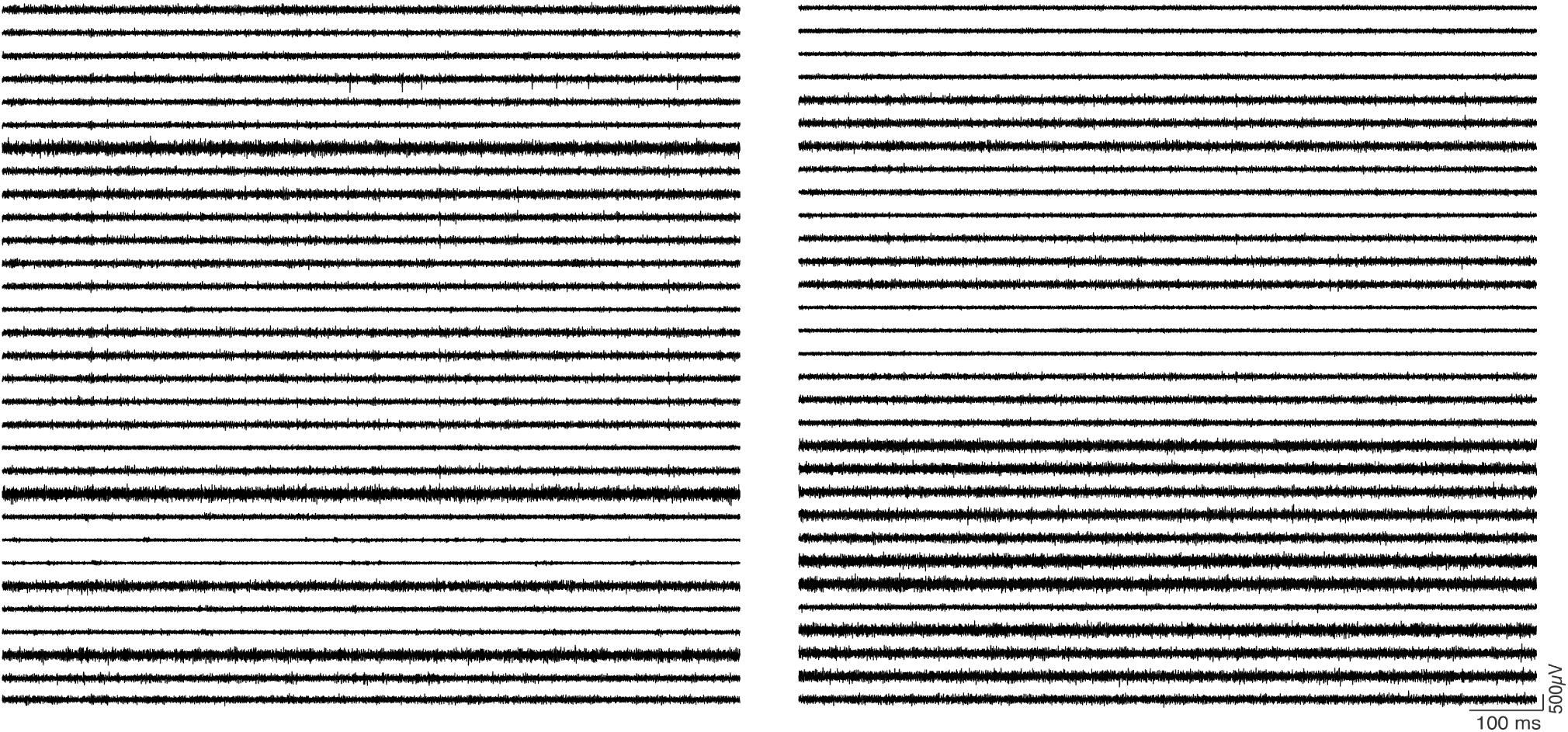
One second of raw filtered activity recorded from a 64-channel CF array implanted in rat V1 showing multiunit signals.

**Figure 6.**
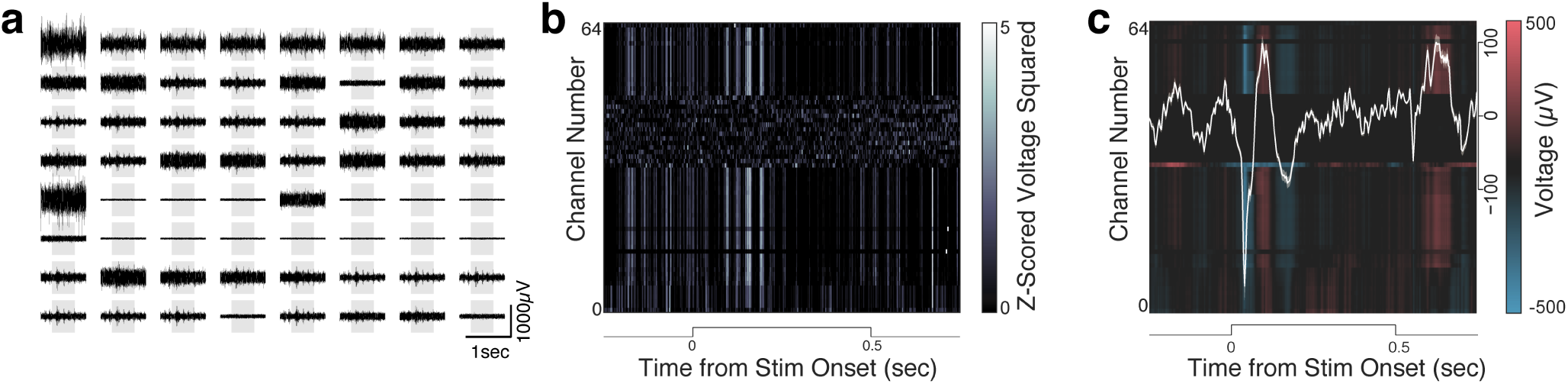
Visual cortex responses to visual stimulation. The stimulus was a 500-millisecond light flash delivered while the rat moved freely in the recording chamber. (**a**) Filtered raw traces across 64 CF electrodes during one stimulation trial (grey patch symbolizes the duration of the light flash) showing multiunit (MU) responses to the stimulus. (**b**) Mean MU response across 225 trials of the light flash (filtered traces were z-scored, squared, and smoothed with a gaussian filter). (**c**) Mean LFP (300 Hz low-pass filtered) response across the same trials (white line: mean LFP response across channels).

## Discussion

CFEAs have shown promise in recording from neuronal populations over long timescales but have so far been prepared in small channel-count configurations^25, 27, 34–36^. It is becoming increasingly clear that in order to understand the brain, neuroscientists will need to measure larger populations of neurons spread out across multiple brain regions^6^ (however, see Gao et. al^37^). Our results show the feasibility of producing high channel-count, high-density CFEAs and using them to record evoked activity in rat visual cortex. Further, we have shared designs to implement simultaneous recordings from multiple brain regions using four 16-channel flex-PCB CFEAs (Fig. 1c) and micro-drive-enabled 64-channel arrays (Fig. 1d). We have also designed a jig for setting up Parylene-C deposition onto the bare CFs (Fig. 3), which we hope will facilitate the implementation of CFEAs by other labs.

The 64-channel CFEAs are faster to assemble (per channel) than their 16-channel predecessors (∼2 hours for the former and ∼1 hour for the latter: a 2X improvement from 3.75 to 1.875 minutes/channel). The method of preparing the recording sites using acid etching and electroplating with PEDOT-TFB further sped up the assembly process over the previous fire-sharpening technique, which was done on individual fibers and required extensive practice^27^.

CFEA designs have for the most part required manual assembly. This will obviously make recording extremely large populations of neurons (10^5-6^) a challenge. As such, manually-assembled CFEAs may be a more appropriate replacement for other manually-made electrodes like tetrodes, rather than arrays manufactured using nanotechnology^6, 22, 23^, which may soon be able to accommodate thousands of recording sites. Still, CFEAs are smaller and more flexible than most fabricated arrays and produce a smaller immune response than larger implants^9^. CFEAs can, in principle, be manufactured using automated systems, and some efforts are underway to implement those^38^.

Electrode arrays are currently the best tool for investigating neuronal circuits continuously (i.e. 24/7) over long periods of time in freely behaving animals. However, whether electrode arrays of any kind will be the method to record all the neuronal activity in a brain is still unknown. Arrays designed to capture the entirety of neuronal activity in a vertebrate brain would need to be quite noninvasive indeed; they would also need to capture the activity of millions of neurons, which would require multiplexing and digitization near the recording sites^39^.

Alternatives to electrodes might be optical methods, which are currently limited in their ability to image deep brain structures^40^. Imaging of calcium indicators has a limited temporal resolution because of molecular binding kinetics^41^; this problem may be avoided by voltage sensors, but those indicators currently have low signal quality^42, 43^ (but see Adam et. al^44^, and in principle suffer from the same depth imaging and bleaching problems as calcium indicators. One radically different approach is molecular barcoding, in which neuronal activity is recorded by a ‘molecular ticker-tape’ that tracks changes in intracellular calcium concentration by making insertions into a cell’s DNA^45, 46^. For now, perhaps the best way to get around the limitations of each technique is to use multiple complementary tools, investigating local circuits with electrode arrays after a brain region of interest is identified using whole-brain low-temporal-resolution methods like calcium imaging or fMRI.

## Methods

### Animals

The care and experimental manipulation of all animals were reviewed and approved by the Harvard Institutional Animal Care and Use Committee (Protocol #27-22). Experimental subjects were female Long Evans rats 3 months or older, weighing 300-500 grams (*n* = 3, Charles River). One rat was implanted with a 16-channel array; two were implanted with 64-channel arrays.

### Carbon Fibers, Array Assembly, and Tip Preparation

In this report, we used 4.5*µ*m diameter carbon fibers (UMS2526, Goodfellow USA), as reported previously^27^. Epoxy sizing was removed either by heating the fibers to 400 C in a kiln, or by soaking the fibers in acetone >24 hours^47^. To insulate the fibers with Parylene-C (di-chloro-di-p-xylylene) (Paratronix Inc, Westborough MA), we laid the fibers out onto sticky notes attached to plastic cartridges (Fig. 3) and coated the fibers with ∼4.5*µ*m layer of Parylene-C using chemical vapor deposition in a PDS-2010 Labcoter machine (Specialty Coating Systems, http://scscoatings.com/), as described previously^27^.

The fibers were then threaded through 3D-printed plastic blocks, which were custom-designed in SolidWorks (Dassault Systèmes SolidWorks Corporation, Waltham, MA) and manufactured using stereolithography (Realize Inc, Noblesville, IN) (designs available on https://github.com/guitchounts/electrodes). The fibers were left protruding ∼1mm from the top of the plastic block and were then deinsulated with a lighter flame, after which they were threaded further into the block until no longer protruding. Connectorization was achieved by flooding the block’s wells with silver paint (MG Chemicals, #842-20g) and attaching a Hirose DF40 connector (Digi-key, H11630CT-ND).

The recording sites were prepared by cutting the array bundles to the desired length for implantation (∼1mm for the visual cortex implants described here) using serrated scissors (Fine Science Tools, 14058-11), then dipping the tips into 99% sulfuric acid (Sigma, 339741-500ML) for 5 minutes, then (after washing any remaining acid off with water), by electroplating with PEDOT-TFB^30, 31^ by applying a 2nA current via a 10V power supply in series with a 5000MΩ resistor (Digikey, SM102035007FE-ND), for 1 minute (for a total charge delivery of 120nC) per electrode^25^. The electroplating solution consisted of 0.01M EDOT (Sigma, 483028-10G) and 0.1M tetrabutylammonium tetrafluoroborate (TFB) (Sigma, 86896-25G) in acetonitrile (Sigma, 360457-500ML). The impedance was tested at 1kHz using an Intan-based headstage and OpenEphys GUI (http://www.open-ephys.org/gui/).

### Surgery

Animals were anesthetized with 2% isoflurane and placed into a stereotaxic apparatus (Knopf Instruments). Care was taken to clean the scalp with Povidone-iodine swabsticks (Professional Disposables International, #S41125) and isopropyl alcohol (Dynarex #1204) before removing the scalp and cleaning the skull surface with hydrogen peroxide (Swan) and a mixture of citric acid (10%) and ferric chloride (3%) (Parkell #S393). Three skull screws (Fine Science Tools, #19010-00) were screwed into the skull to anchor the implant. A 0.003” stainless steel (A-M Systems, #794700) ground wire was inserted ∼2mm tangential to the brain over the cerebellum.

The arrays were targeted to right V1, ranging ∼6-8 mm posterior to bregma, 4.5 mm ML, reaching layer 2/3 at 0.6 mm DV. After electrodes were inserted into the brain, the craniotomy was covered with Puralube vet ointment (Dechra) and the electrodes were glued down with Metabond (Parkell). Post-operative care included twice-daily injections of buprenex (0.05 mg/kg Intraperitoneal (IP)) and dexamethasone (0.5 mg/kg IP) for three days.

### Electrophysiology

Electrode signals were acquired at 30 kHz using either a 16-channel Intan RHD2132 headstage, or a custom Intan-based 64-channel headstage^48^ and Opal-Kelly FPGAs (XEM6010 with Xilinx Spartan-6 ICs). LFPs and spikes were extracted following procedures described in Dhawale et al^48^. The LFP signals were downsampled to 300Hz by two applications of a fourth order 5-fold decimating Chebychev filter followed by a single application of a fourth order 4-fold decimating Chebychev filter. The 16-channel data were acquired using the OpenEphys GUI^49^.

In addition to the 64-channel CFEAs, we designed a 64-channel system for recording with four 16-channel arrays in multiple brain regions. The 16-channel flex-PCB (Fig. 1c) mates with a miniature 16-channel 3D-printed plastic block that holds the CFs. The connector side holds a 20-pin Hirose DF40 connector (Digi-key, H11618CT-ND) that interfaces with a breakout PCB that holds four mating 20-pin connectors (Digi-key, H11619CT-ND) on one side and a 70-pin DF40 connector (Digi-key, H11630CT-ND) on the other. The breakout board thus connects four flex-PCB arrays to one 64-channel headstage. The 64-channel CFEAs were also adapted for microdrive-based recordings based on the optetrode design in Anikeeva et. al^32^ (Fig. 1 d).

### Behavior and Visual Stimulation

The recordings were carried out in a custom recording chamber while the animals were free to behave. The headstage was connected to the FPGA board via a custom shielded cable (Samtec, SFSD-07-30C-H-12.00-DR-NDS, TFM-107-02-L-D-WT; McMaster extension spring 9640K123) and commutator (Logisaf 22mm 300Rpm 24 Circuits Capsule Slip Ring 2A 240V Test Equipment, Amazon).

Visual stimulation was delivered via white LED lights (Triangle Bulbs Cool White LED Waterproof Flexible Strip Light, T93007-1, Amazon) mounted in the recording chamber, and controlled with a micro-controller (Arduino). During stimulus presentation, the chamber and experimental room were otherwise completely dark. The lights were On for 500 ms and Off for a uniform-random time between 400 and 600 ms.

### Imaging

Scanning Electron Microscopy images of the CF tips were taken using a Zeiss Ultra55 Field Emission Scanning Electron Microscope (FESEM) at the Center for Nanoscale Systems at Harvard University.

## Acknowledgements

We would like to thank Andrea Stacy for assistance with SEM imaging at the Center for Nanoscale Systems at Harvard and Guosong Hong for advice on acid etching. We are immensely grateful to Jeffrey Markowitz, Steffen Wolff, Robert Johnson, and Javier Masis for comments on the manuscript. GG was supported by the National Science Foundation (NSF) Graduate Research Fellowship Program (GRFP).

## Author contributions statement

GG conceived and performed the experiments, analyzed the data, and wrote the manuscript. DC provided research funding.

